# Spider mites escape bacterial infection by avoiding contaminated food

**DOI:** 10.1101/408203

**Authors:** Flore Zélé, Gonçalo Santos-Matos, Alexandre Figueiredo, Cátia Eira, Catarina Pinto, Telma Laurentino, Élio Sucena, Sara Magalhães

**Affiliations:** Centre for Ecology, Evolution and Environmental Changes (cE3c), Faculdade de Ciências da Universidade de Lisboa, Edificio C2, 3° Piso Campo Grande, 1749-016 Lisboa, Portugal; Instituto Gulbenkian de Ciência, Apartado 14, 2781-901 Oeiras, Portugal; Department for Plant and Microbial Biology, University of Zurich, Winterthurerstrasse 190, 8057 Zürich, Switzerland; Department of Evolutionary Biology and Environmental Studies, University of Zurich, Winterthurerstrasse 190, 8057 Zürich, Switzerland; Zoological Institute, University of Basel, Vesalgasse 1, 4051 Basel, Switzerland; Departamento de Biologia Animal, Faculdade de Ciências da Universidade de Lisboa, Campo Grande, Edifício C2, 1749-016 Lisboa, Portugal

**Keywords:** Parasitism, immunity, behavioural avoidance, *Tetranychus urticae*

## Abstract

To fight infection, arthropods rely on the deployment of an innate immune response but also upon physical/chemical barriers and avoidance behaviours. However, most studies focus on immunity, with other defensive mechanisms being relatively overlooked.

We have previously shown that the spider mite *Tetranychus urticae* does not mount an induced immune response towards systemic bacterial infections, entailing very high mortality rates. Therefore, we hypothesized that other defence mechanisms may be operating to minimize infection risk. Here, we test (a) if spider mites are also highly susceptible to other infection routes - spraying and feeding - and (b) if they display avoidance behaviours towards infected food. Individuals sprayed with or fed on *Escherichia coli* or *Pseudomonas putida* survived less than the control, pointing to a deficient capacity of the gut epithelium, and possibly of the cuticle, to contain bacteria. Additionally, we found that spider mites prefer uninfected food to food contaminated with bacteria, a choice that probably does not rely on olfactory cues.

Our results suggest that spider mites may rely mostly on avoidance behaviours to minimize bacterial infection and highlight the multi-layered nature of immune strategies present in arthropods.

## INTRODUCTION

Arthropods are often attacked by pathogens, leading to severe fitness costs. In response, arthropods have evolved several mechanisms to counter infection, ranging from behavioural avoidance to the deployment of an immune response. While a large body of literature concerns the immune response of arthropods to bacterial infection (Schmid-Hempel 2005; Lemaitre and Hoffmann 2007; Graham et al. 2011), these were found to be inefficient in fighting infection in both aphids (Altincicek et al. 2008; Gerardo et al. 2010; but see Laughton et al. 2016; Parker et al. 2017) and spider mites (Santos-Matos et al. 2017) compared to other invertebrate studied to date.

Systemic immunity operates once a pathogen is inside the body of a host. However, pathogens must cross a series of barriers before entering their hosts (Hall et al. 2017). Indeed, many pathogens are ingested by their host but need to bypass the gut epithelium to cause infection, point at which most pathogens are contained (Kuraishi et al. 2011; Martins et al. 2013). Such strategies are not detected by classical immunological studies, which investigate responses upon injection of the parasite into the body cavity of the host. Moreover, ingestion of potential pathogenic agents may be controlled through host behavioural responses (Clayton 1991). However, despite their importance as a first line of defence against parasites (Hart 1990; Hart 1994) and their predicted strong effect on host-parasite dynamics (Eakin et al. 2015), behavioural strategies to avoid infection remain much less studied than ‘classical’ immune responses (reviewed by Parker et al. 2011), or than behavioural strategies to avoid predation (reviewed by Buck et al. 2018). Moreover, avoidance behaviour is likely the most cost-effective strategy employed by free-living animals to maintain fitness in the face of parasite threat, as compared to resistance and tolerance (Curtis 2014). Indeed, many behavioral strategies allowing animals to avoid exposure to parasites, and thus the likelihood of infection, have been described (for reviews, see Buck et al. 2018; Cremer et al. 2018; Curtis 2014; Moore 2002). For example, hosts may detect parasites and avoid them directly, such as aphids dropping from plants to escape parasitoid wasps (Fill et al. 2012); or avoid habitats harbouring pathogens, such as the rainbow trout avoiding shelters where eye fluke cercariae are present (Karvonen et al. 2004). Hosts may also avoid contact with infected conspecifics, such as bullfrog tadpoles when facing yeast infection (Kiesecker et al. 1999). Finally, hosts may avoid ingesting infected food, such as gypsy moth larvae that avoid feeding on foliage contaminated with cadavers of virus-infected conspecific larvae (Parker et al. 2010). Some organisms are also able to avoid different sources of infection, such as adult seven-spot ladybirds, which avoid the fungus *Beauveria bassiana* on leaf surfaces, in soil and in mycosed conspecifics cadavers (Ormond et al. 2011). However, some organisms lack pathogen avoidance behaviours (Mnyone et al. 2010); or may even be attracted toward infected conspecifics (Bouwman and Hawley 2010; Cornet et al. 2013).

We have previously shown that most genes acting in *Drosophila* immunity, which are generally conserved among arthropods, are absent in the spider mite *Tetranychus urticae* (Grbic et al. 2011). Furthermore, we showed that spider mites do not mount an immune response and die upon infection by both gram-positive and gram-negative bacteria (Santos-Matos et al. 2017). However, these conclusions were based upon experiments using bacterial injection, a route of infection that bypasses a series of possible barriers to infection. Here, we set out to investigate (a) whether spider mites are also susceptible to bacterial infection by spraying (i.e., topical infections and/or bacteria ingestion) or to bacteria ingestion only, (b) whether they are able to avoid contaminated food, and (c) whether olfactory cues play a role in such avoidance. To this aim, we used two species of bacteria: a non-pathogenic strain of *Escherichia coli,* commonly used in immunity studies (Lemaitre et al. 1996; Santos-Matos et al. 2017) and found on plants (Meric et al. 2013; Seo and Matthews 2012; Solomon et al. 2003; Solomon et al. 2002); and *Pseudomonas putida,* a bacterium commonly found on plants (Bodenhausen et al. 2013; Rastogi et al. 2012), which is pathogenic to spider mites (Aksoy et al. 2008) and other arthropods (Ateyyat et al. 2010; Ayres and Schneider 2008; Schneider and Dorn 2001) as other *Pseudomonas* species (Broadbent and Matteoni 1990; Bucher and Stephens 1957; Commare et al. 2002; Poinar and Poinar 1998; Thomas and Poinar 1973; Vodovar et al. 2006; Vodovar et al. 2005).

## Material and Methods

### Spider mite populations

All experiments were done with the two-spotted spider mite, *Tetranychus urticae.* For the olfactory cue experiments, we used the TuTom.tet population, originally collected from tomato plants in Carregado, Portugal, in August 2010 (Clemente et al. 2016). Other experiments were performed using a population derived from the London strain. This strain was originally collected from fields in the Vineland region, Ontario, Canada and was used to sequence the species genome (Grbic et al. 2011). Both populations were maintained under controlled conditions (25 ± 2°C, 16/8 h L/D) on bean plants (*Phaseolus vulgaris,* variety Enana, provided by Germisem, Portugal) at the Faculty of Sciences of the University of Lisbon, since 2010. To control for age of the tested females, adult spider mite females were placed on a bean leaf inside a petri dish, where they laid eggs for 24 hours. These petri dishes were then kept under the aforementioned controlled conditions and spider mites developed into adulthood. For the experiments, we used 0-3 day-old females (since the last moult).

### Bacterial stocks

Bacterial stocks of *E. coli* DH5*α* (Gram negative) and *P. putida* (Gram negative) were retrieved from a -80°C stock and plated on Petri dishes with Luria Broth (LB) 3 days before each experiment was performed. Subsequently, a colony was picked from the Petri dishes, transferred to liquid LB, grown overnight at 37° for *E. coli* and 30° for *P. putida* and diluted in LB (i.e. the same LB solution than the control: 10g/L tryptone; 5g/L Yeast Extract; 10g/L NaCl) to the required concentrations: optical density (OD) of 1, 10 and/or 25, measured at 600nm with GENESYS™ 10S UV spectrophotometer (Thermo Scientific™), depending on the experiment. Although OD is a standard measurement in microbiology, the concentration of both bacteria species may differ for a same OD value (e.g. OD1 of *E. coli* corresponds to 8 x 10^8^ cells/ml, while OD1 of *P. aeruginosa* correspond to 2 x 10^8^ cells/mL; Kim et al. 2012). Since the concentration of *P. putida* is, to our knowledge, unknown, we performed a simple test to determine its concentration relative to that of *E. coli.* 100 μl of solution of each species (bacteria at OD1 serially diluted to 1:800000) were plated in 5 different petri dishes, and the number of Colony Forming Units (CFUs) was counted in each dish the next day. Plates inoculated with *P. putida* contained 57.8 ± 3.2 CFUs, while those with *E coli* had 145.4 ±6.4 CFUs.

### Effects of bacteria spraying on spider-mite mortality and oviposition

The effects of *E. coli* and of *P. putida* infection by spraying were tested in two separate experiments, each consisting of 3 independent experimental blocks. For each block, 200 *T. urticae* females were individually placed on bean leaf fragments (1.5cm length and 1cm width) placed on wet cotton inside Petri dishes (up to 25 leaf fragments per Petri dish). This procedure prevents spider mites from escaping the leaf fragments (individuals that fall out of the leaf fragments are drowned in the wet cotton). Subsequently, each Petri dish was sprayed three times using a sprinkler (0.33 ml per spatter; c.a. 1ml per Petri dish), at a height of 30cm, with bacteria at OD1, ODIO, or OD25, or with LB as control (i.e., 50 females were allocated to each treatment). Subsequently, spider mites were kept in a controlled environment (25 ± 2°C, 16/8 h L/D) for 96 hours. Survival and fecundity were recorded every 24 hours. For each bacterial species, a total of 600 mites were thus tested (50 per treatment per block).

### Effects of bacteria ingestion on spider-mite mortality and fecundity

To measure spider mite mortality and fecundity upon bacterial ingestion, 40 females were placed on circular cardboard arenas (ca. 3cm^2^) with 2 parafilm bubbles (made using a vacuum manifold attached to a vacuum pump and subsequently closed using adhesive tape). The bubbles were filled with 30 |il of food consisting of a 1:1 proportion of Schneider’s medium and either LB (negative control) or LB with bacteria at OD1, ODIO, or OD25. A green food colouring dye was added to the mixture at a 1:4 proportion to distinguish spider mites that fed on the bubbles from those that did not (mites are relatively transparent hence the dye is easily seen inside mites). Mites were then kept in the arenas for 48 hours before scoring the number of females alive and those with dye in their gut. Females that had dye in the gut were individually placed on bean leaf fragments (1.5cm length and 1cm width) over wet cotton inside Petri dishes (up to 25 leaf fragments per Petri dish) and kept in a controlled environment (25 ± 2°C, 16/8 h L/D) for 96 hours during which survival and fecundity were recorded every 24 hours. As mentioned above, the effects of infection by *E. coli* and *P. putida* were tested in two separate experiments with an initial number of 400 mites each and consisting of 5 independent experimental blocks, with two arenas per block.

### Effects of volatile cues on spider mite avoidance behaviour

To investigate whether spider mites use olfactory cues to avoid food sources contaminated with bacteria, we performed two experiments using previously established protocols (Pallini et al. 1997; Rodrigues et al. 2017). In a first experiment, we performed a two-way choice test where adult females were placed in the middle of a dumbbell-shaped arena consisting of a strip of parafilm (2 cm x 0.5 cm) connecting one bean leaf disc (0.64 cm^2^) with a droplet (10 μ.L) of LB and another with either no droplet (control), a droplet containing *E. coli* (ODIO), or a droplet containing *P. putida* (ODIO). Trials began as soon as individuals were placed in the centre of the parafilm bridge and lasted 30 minutes. The time spent by a female on either side of the arena was recorded. A test was considered invalid if an individual drowned or failed to leave the centre of the arena. A total of 66 females were tested in this experiment (n=21 for the control, n=22 for *P. putida,* and n=23 for *E. coli).*

In a second experiment, we used a Y-maze olfactometer connected to a vacuum pump with an airflow of 0.4-0.5 m/s to test avoidance off. *coli* (OD25) or of *P. putida* (OD25) by *T. urticae* females. Two types of tests were performed. In the first, we tested whether spider mites use olfactory cues to avoid bacteria in absence of the food source. Cotton soaked in LB was placed inside a syringe connected to one of the Y-maze arm, while cotton soaked in bacteria was connected to the other. In the second test, we tested whether spider mites use olfactory cues to avoid food sources contaminated with bacteria. The day before the test, very early in the morning (optimal for pathogen proliferation), bean plants (14 days-old) were covered with a transparent plastic dome inside the growing chamber (25°C, 16/8 h L/D) and soaked in water for 2-4 hours to increase stomatal opening (Liu et al. 2015). 24h before the test, 24 droplets of 10μl of the bacterial suspension at OD25 or of LB (as control), were placed on two leaves of the experimental plants. The plants were then placed under a transparent plastic dome under the light in the experimental room for one day to maintain humidity (Liu et al. 2015) and to avoid communication between plants. On the day of the test, both bean plants with LB or with bacteria were placed individually in an experimental plastic box (18.5x18.8x24.5cm) connected to each arm of the Y-maze. For both tests, PES Sterile Membrane Filters (0.2μm; VWR™) were connected to the tip of the tube connecting the syringes or the plants’ boxes to the olfactometer to ensure that bacteria did not contact with the olfactometer arms and that only olfactory cues could be perceived. One-day old adult females were then placed at the end of the Y-maze and allowed to move along the central wire of the olfactometer, without touching the glass walls. Each female was tested individually and a choice was considered valid when the mite entered the last 1/3 of one of the olfactometer arms within five minutes. To avoid confounding effects, bacteria were swapped from one olfactometer arm to the other every 5 valid choices. For each test, a total of 80 valid choices were obtained within four experimental blocks (20 choices per block), each experimental block corresponding to a different day.

### Statistical analyses

Analyses were carried out using the R statistical package (v. 3.3.2). The different statistical models built to analyse the effect of bacterial infection on the survival, fecundity and behaviour of *T. urticae* spider mites are described in the Table S1 of the electronic supplementary material (Additional file 1).

To analyse the effect of bacteria on survival, oviposition and feeding behaviour of spider mites (models 1 to 10), the general procedure to build the statistical models was as follows: the treatment (bacterial concentration to which mites were exposed: LB only [i.e., OD0 for bacteria], OD1, ODIO, OD25) was fit as a fixed explanatory variable, whereas blocks were fit as random explanatory variables. Survival data (models 1, 3, 5, 7) were analysed using a Cox proportional hazards mixed-effect model (coxme, kinship package). Hazard ratios were obtained as an estimate of the difference in survival rates (Crawley 2007) between the control (LB) and each of the other factor levels. As the oviposition data (number of eggs laid per female per day) were greatly overdispersed, they were analysed using a mixed model gimmadrmb procedure (glmmADMB package). For these models (models 2, 4, 6, 8), error structures that best fit the errors distribution (e.g. zero-inflated quasi-poisson or negative binomial) were chosen based on AIC, the variable “day” was added as fixed explanatory variable and mite identity nested within blocks was fit as random explanatory variable across “days” to account for temporal autocorrelation across repeated measures on the same individuals (Crawley 2007). When a significant interaction was found between the variables “day” and “treatment”, the effect of treatment was analysed within each day separately. The proportion of mites that fed (models 9 and 10) was computed using the function cbind (number of females fed *vs.* unfed) and subsequently analysed either with a mixed model glmmadmb procedure (glmmADMB package) with zero-inflated binomial error distribution to account for overdispersion (model 9), or with a generalized linear mixed effect model (glmer, lme4 package) with a binomial error distribution (model 10). In all analyses, when the variable “treatment” was significant, a stepwise *a posteriori* procedure (Crawley 2007) to determine differences between treatments was carried out by aggregating factor levels together and by testing the fit of the simplified model using a likelihood ratio test (LRT), which is approximately distributed as a χ^2^ distribution (Bolker 2008).

To analyse the effect of olfactory cues on spider-mite avoidance of contaminated food (model 11), the time spent by each female on each side of the arena was computed using the function cbind (time spent on the side of the droplet containing bacteria or no droplet *vs.* time spent on the side of the droplet containing LB), and subsequently analysed using a generalized linear model (glm) using a quasibinomial error distribution to account for over-dispersion. The type of test (LB vs no droplet, a droplet with *E. coli,* or with *P. putida)* was fit as fixed explanatory variable along with the side (left or right) on which the control (LB) droplet was located. To determine whether, in each treatment, the mites spent significantly more time in one of the sides of the arena, the intercept of the model was changed to zero, which gives (in a model with categorical factors and a binomial distribution) the estimates of each fixed factor against a probability of 0.5 (Crawley 2007).

To analyse the effect of olfactory cues on spider-mite avoidance behaviour using the olfactometer (model 12), the type of test (LB vs *E. coli* or *P. putida* on cotton or on the entire plant) was fit as fixed explanatory variable, whereas block and the side (left or right) on which the control (LB) was located were fit as random explanatory variables. The olfactometer arm chosen by mites, a binary response variable (0: LB 1: bacteria) was then analysed using a generalized linear mixed effect model (glmer, lme4 package) with a binomial error distribution. As before, the intercept of the model was changed to zero to determine whether the probability of a mite choosing one arm of the olfactometer differs significantly from 0.5.

For all analyses, maximal models were simplified by sequentially eliminating nonsignificant terms to establish a minimal model (Crawley 2007), and the significance of the explanatory variables was established using χ^2^-tests or *F-tests* to account for over-dispersion (Bolker 2008). The significant values given in the text are for the minimal model, while nonsignificant values are those obtained before removing the variable from the model (Crawley 2007). Full datasets are given in Additional files 2 to 7.

## RESULTS

### Effects of bacteria spraying on spider-mite mortality and fecundity

Spraying of *T. urticae* females with *E. coli* negatively affected their survival (treatment effect: *X*^*2*^_*3*_ *-* 80.21, *P* < 0.0001; model 1; Figure 1A). Contrast analyses revealed that mortality of mites sprayed with bacteria at ODIO did not increase compared to those sprayed with a concentration of OD1 (contrasts OD1 us. ODIO: *X*^2^ _*3*_ = 1.01, *P* = 0.32). However, mortality caused by these treatments was 8.6-fold higher than that of mites sprayed with LB only (contrasts ODI-ODIO *vs.* LB: *X%* = 104.02, *P* < 0.0001), and 4.12-fold lower than that of mites sprayed with *E. coli* at OD25 (contrasts ODI-ODIO us. OD25: *X*^*2*^ _*1*_ = 9.13, *P* = 0.003). Spider mite oviposition increased similarly through time in all treatments (day*treatment interaction: *X*^*2*^_*3*_ = 6.82, *P* = 0.08; Main effect of day: *X*^*2*^ _*3*_ = 15.76, *P <* 0.0001; model 2; Figure 1B) but differ significantly between treatments (Main effect of treatment: *X*^*2*^_*3*_ *=* 35.80, *P <* 0.0001; model 2; Figure 1B). Indeed, contrast analyses showed that, although *E. coli* at OD25 did not significantly affect oviposition (contrasts LB *vs.* OD25: *X*^*2*^ _*1*_ = 1.40, *P* = 0.24), females laid more eggs per day after being sprayed with *E. coli* at OD1 or ODIO (contrasts OD1 *vs.* ODIO: *X*^*2*^ _*1*_ *=* 0.14, *P* = 0.71; LB-OD25 us. ODI-ODIO: *X*^*2*^_*1*_ = 34.36, *P* < 0.0001).

**Figure 1.**
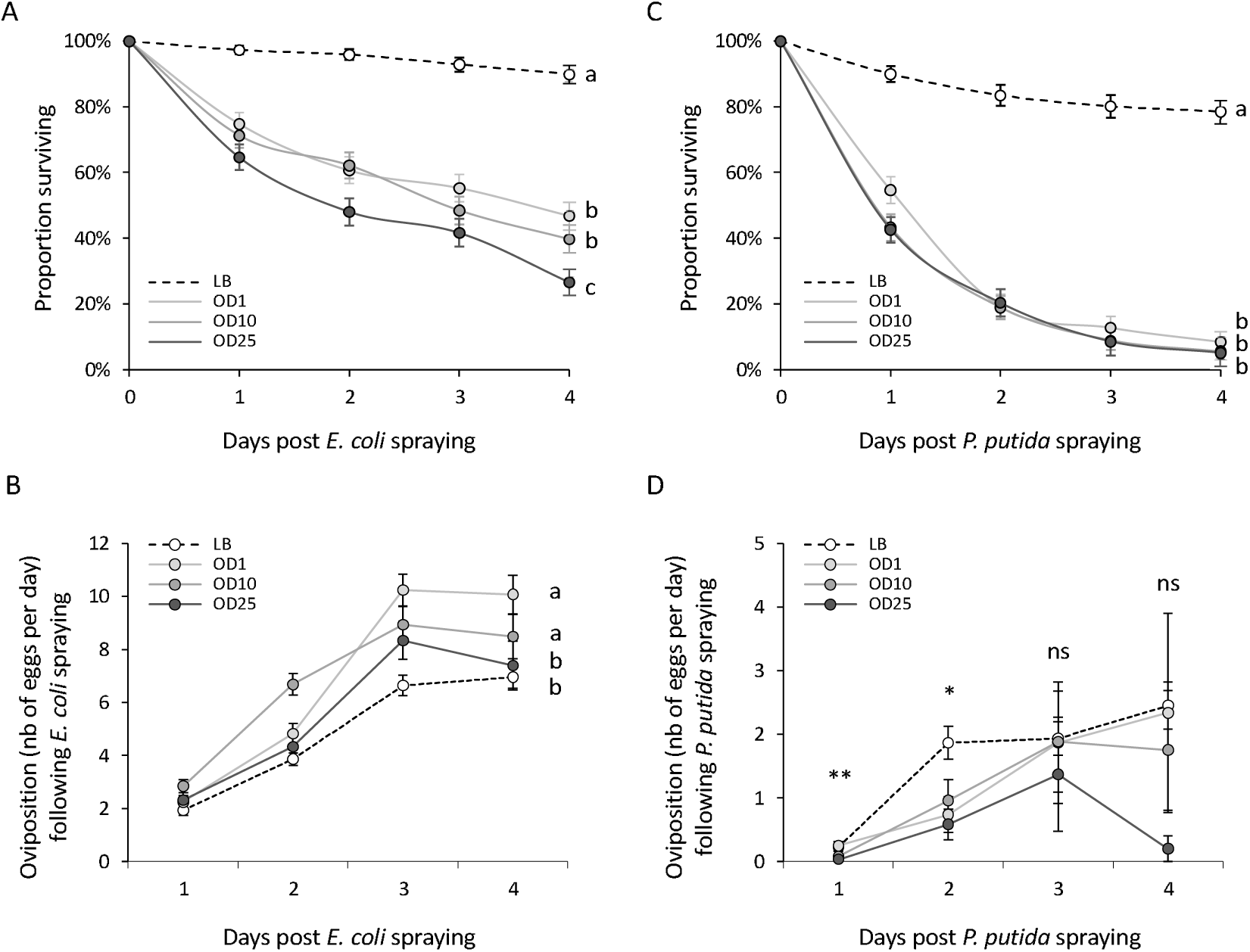
Effect of bacteria spraying on spider-mite survival (A, C) and daily oviposition (B, D). Females were exposed to different concentrations of either *E. coli* (A, B) or *P. putida* (C, D). In (B) and (D), points represent the mean number of eggs (± s.e.) for each experimental day. In (A), (B) and (C), identical letter superscripts indicate non-significant differences between treatments at the 5% level (*a posteriori* contrasts). In (D), as a significant interaction between day and treatment was found, superscripts indicate the significance of the treatment effect for each day (ns: no significant differences at the 5% level, *p < 0.05, ** p< 0.01). White dots and black dashed line: LB; Light grey dots and lines: OD1; Medium grey dots and lines: ODIO; Dark grey points and lines: OD25.

Similarly to *E. coli,* spraying of *P. putida* decreased female survival (main effect of treatment: *X*^*z*^_*3*_ *=* 138.19, *P <* 0.0001; model 3; Figure 1C) to the same extent for all bacterial concentration used (contrasts OD1 us. ODIO us. OD25: *X*^*2*^_*2*_ *-* 2.58, *P* = 0.28). Contrasting with the results for *E. coli,* the effect of *P. putida* on spider-mite oviposition varied with the number of days after spraying (day*treatment interaction: *X*^*2*^_*3*_ = 8.12, *P* = 0.04; model 4; Figure 1D). Indeed, the separate analyses of each day following spraying revealed that *P. putida* decreased significantly the oviposition of spider mites during the first two days following spraying (main effect of treatment at day 1: *X*^*2*^_*3*_ *=* 15.10, *P* = 0.002, and at day 2: *X*^*2*^_*3*_ *=* 9.51, *P* = 0.02). During the first day females sprayed with ODIO or OD25 (contrasts ODIO us. OD25: *X*^*2*^*i* = 1.66, *P* = 0.20) laid fewer eggs than those sprayed with LB or OD1 (contrasts LB us. OD1: *X*^*2*^*!* = 0.31, *P* = 0.58; LB-OD1 us. OD10-OD25: *X*^*2*^ _*3*_ *=* 13.14, *P <* 0.001), while during the second day all bacterial doses led to decreased oviposition compared to the control LB to a similar extent (contrasts OD1 us. ODIO *vs*. OD25: *X*^*2*^_*2*_ *-* 0.18, *P* = 0.91). In contrast, on the following two days no effect on oviposition could be detected (main effect of treatment at day 3: *X*^*2*^ _*3*_ = 0.91, *P* = 0.82, and at day 4: *X*^*2*^_*3*_ = 3.97, *P* = 0.28). Note, however, that for unknown reasons, oviposition was extremely low in the *P. putida* spraying experiment (compare Figures IB and ID).

### Effects of bacteria ingestion on spider-mite mortality and oviposition

Feeding on *E. coli* increased the mortality of *T. urticae* by 4-fold (main effect of treatment: *X*^*2*^_*3*_ *=* 10.36, *P* = 0.02; model 5; Figure 2A), independently of the ingested bacterial concentration (contrasts OD1 *vs.* ODIO *vs.* OD25: *X*^*2*^ _*2*_ = 1.56, *P* = 0.46). The effect of E. *coli* ingestion on spider-mite oviposition varied with the number of days after feeding (day*treatment interaction: *X* _*3*_ *=* 8.10, *P* = 0.04; model 6; Figure 2B). Indeed, spider mite oviposition differed between treatments at days 1 and 2 (main effect of treatment at day 1: *X*^*2*^*3* =17.40, *P <* 0.001, and at day 2: *X*^*2*^_*3*_ *=* 12.85, *P* = 0.005) but not at days 3 and 4 (main effect of treatment at day 3: *X*^*2*^_*3*_ = 3.90, *P* = 0.27, and at day 4: *X*^*2*^_*3*_ = 2.08, *P* = 0.56). For both day 1 and day 2, the females fed on ODIO and OD25 had a decreased oviposition (contrasts ODIO *vs.* OD25 for day 1: *X*^*2*^*i* = 1.74, *P* = 0.19, and for day 2: X^2^_2_ = 0.75, *P* = 0.39) while those fed on OD1 had the same oviposition than those fed on LB only (contrasts LB *vs.* OD1 for day 1: *X*^*2*^ _*1*_ *=* 0.21, *P* = 0.66, and for day 2: *X*^*2*^*!* = 0.04, *P* = 0.84; contrasts LB-OD1 us. OD10-OD25 for day 1: *X*^*2*^ _*1*_ *=* 15.49, *P* < 0.0001, and for day 2: X^2^ _*1*_ = 12.06, *P* = 0.001).

**Figure 2.**
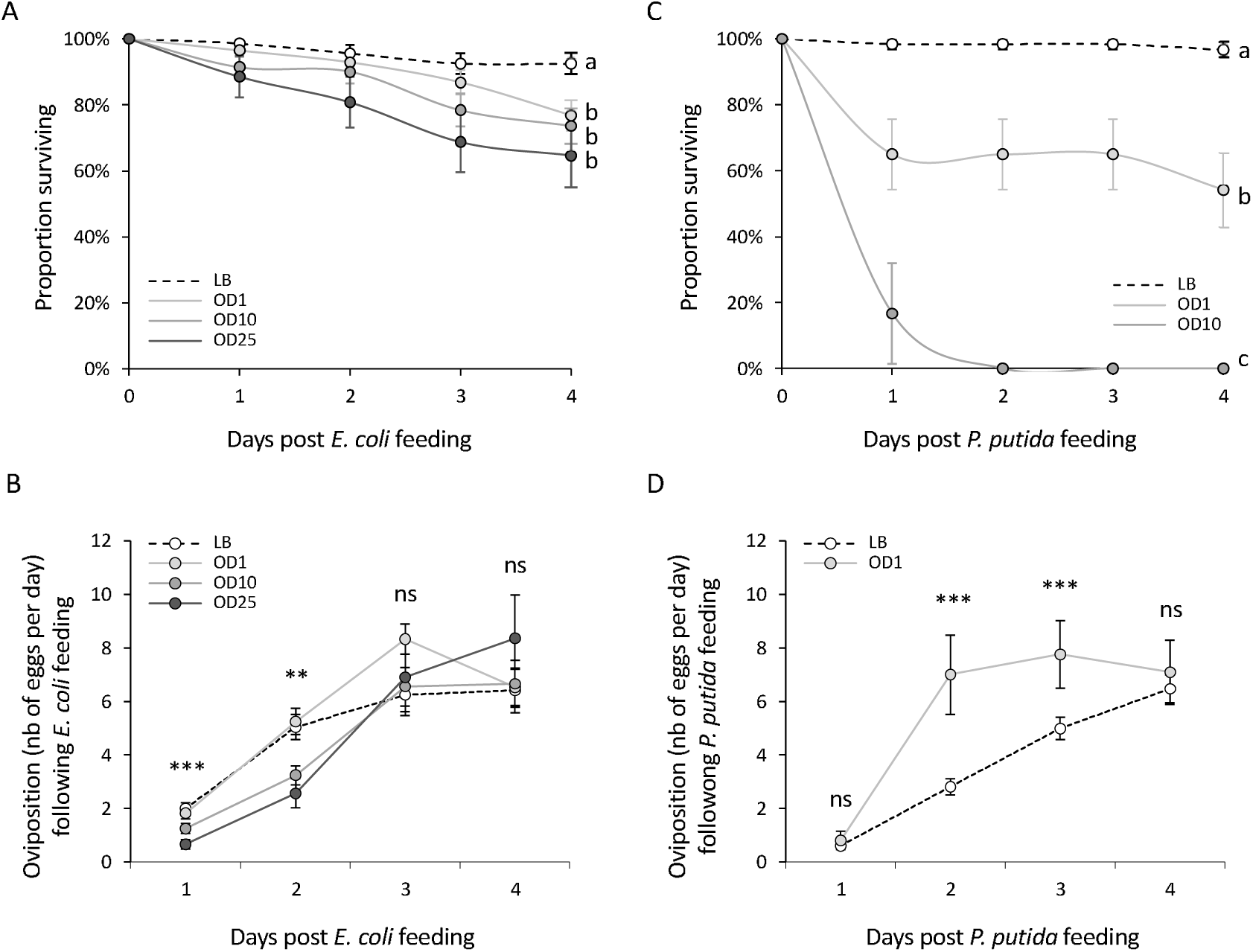
Effect of bacteria ingestion on spider-mite survival (A, C) and daily oviposition (B, D). Females were exposed to different concentrations of either *E. coli* (A, B) or *P. putida* (C, D). In (B) and (D), points represent mean number of eggs (± s.e.) for each day experimental day. In (A) and (C), identical letter superscripts indicate non-significant differences between treatments at the 5% level (*a posteriori* contrasts). In (B) and (D), as significant interactions between day and treatment were found, superscripts indicate the significance of the treatment effect for each day (ns: no significant differences at the *5%* level, **p< 0.01, ***pc 0.001). White points and black dashed line: LB; Light grey dots and lines: OD1; Medium grey dots and lines: ODIO; Dark grey dots and lines: OD25.

As only one mite fed on food contaminated with *P. putida* at OD25 (see below), we could not study the effect of ingestion at this concentration on survival. For the remaining concentrations, feeding on *P. putida* affected spider mite mortality (main effect of treatment: *X*^*2*^ _*3*_ *=* 23.30, *P <* 0.0001; model 7; Figure 2C). Mortality was higher when mites fed on food containing *P. putida* at ODIO than at OD1 (contrasts OD1 *vs.* ODIO: *X*^*2*^ _*1*_ = 4.22, *P* = 0.04), and feeding on bacteria at both concentrations increased spider-mite mortality comparing to control mites (contrasts OD1 *vs.* LB: *X*^*2*^ _*1*_ *=* 19.58, *P <* 0.0001). Since no mites survived after ingestion of *P. putida* at ODIO, only the effect of OD1 on female oviposition could be studied (Figure 2C), and the statistical analyses revealed that this effect varied through time (day*treatment interaction: *X*^*2*^ _*1*_ *=* 6.78, *P* = 0.009; model 8; Figure 2D). While all mites laid the same number of eggs at days 1 and 4 (main effect of treatment at day 1: *X*^*2*^ _*1*_ *=* 0.39, *P* = 0.53, and at day 4: *X*^*2*^ _*1*_ *=* 0.11, *P* = 0.74), at days 2 and 3 the ingestion of *P. putida* at OD1 increased the number of eggs laid compared to the control (main effect of treatment at day 2: *X*^*2*^ _*1*_ *=* 30.34, *P* < 0.0001, and at day 3: *X*^*2*^ _*1*_ *=* 11.01, *P* < 0.001).

### Avoidance of bacteria ingestion

Fewer mites fed on food with *E. coli* than on food with LB only (main effect of treatment: *F*_*3r34*_ *=* 20.13, *P* < 0.0001; model 9; Figure 3A). This effect, however, did not increase with the bacterial concentration, as the proportion of mites avoiding contaminated food was the lowest at ODIO (contrasts LB us. ODIO: *X*^*2*^_*1*_ = 3.85, *P* < 0.05), intermediate at OD1 (contrasts OD1 *vs* ODIO: *X*^2^ _*1*_ = 4.61, *P* = 0.03), and the highest at OD25 (contrasts OD1 *v*s OD25: *X*^*2*^ _*1*_ = 6.69, *P* = 0.01).

**Figure 3.**
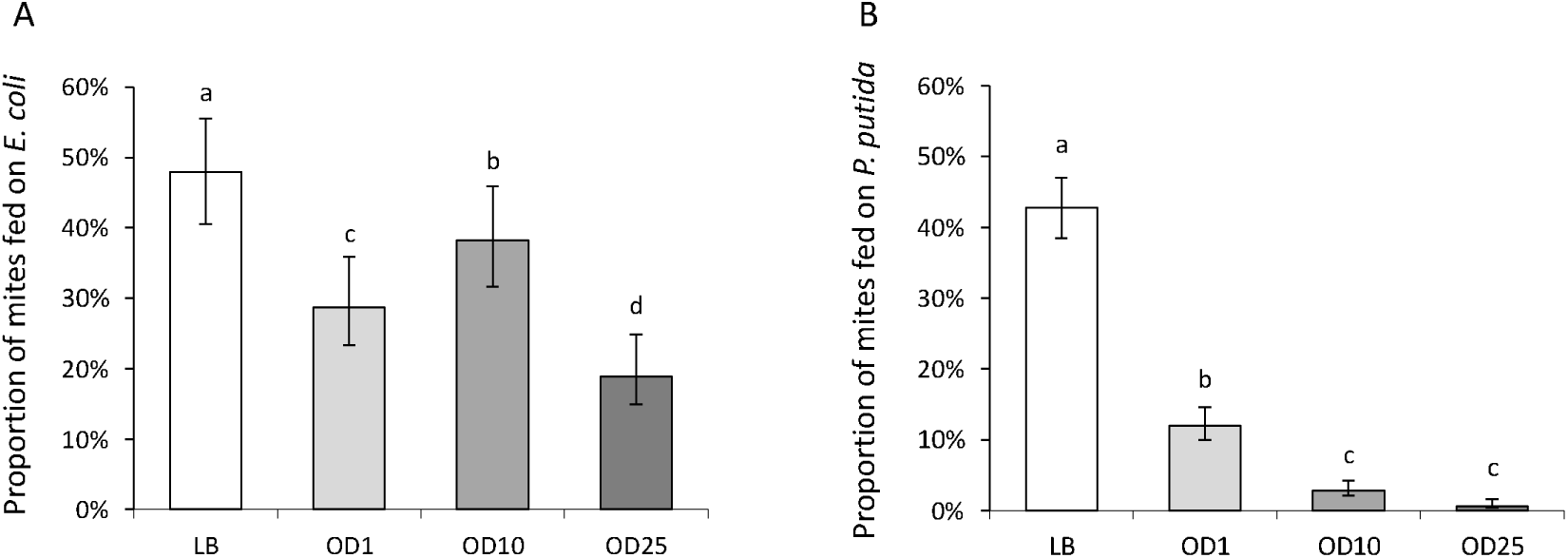
Avoidance of bacteria ingestion by *T. urticae* females. Bars and dots represent mean (± s.e.) proportion of females feeding on *E. coli* (A) or *P. putida* (B). Estimates were obtained from the GLMM statistical mode! that takes into account variation among replicates and blocks as random effect, and corrects for zero-inflation in the case of feeding on *E. coli.* Standard errors were obtained from the upper and lower confidence intervals given by the model. Identical superscripts indicate non-significant differences between treatments at the 5% level (*a posteriori* contrasts). White bars: LB; Light grey bars: OD1; Medium grey bars: ODIO; Dark grey bars: OD25.

Similarly, fewer mites fed on LB with *P. putida* than on LB only (main effect of treatment: *X*^*2*^ _*3*_ = 138.57, *P* < 0.0001; model 10; Figure 3B). Indeed, 43% of the mites fed on LB, but only 12% fed on food containing *P. putida* at OD1 (contrasts LB *vs* OD1: *X*^2^ _*3*_ = 38.62, *P* < 0.0001), and only about 1% fed on food with *P. putida* at ODIO or OD25 (contrasts ODIO vs OD25: *X*^*2*^ _*1*_ = 2.71, *P* = 0.10; contrasts OD1 i/s ODlO-25: *X*^*2*^ _*1*_ = 21.87, *P* < 0.0001).

### Effects of volatile cues on spider-mite avoidance behaviour

When given the choice between leaf discs containing LB with bacteria or LB alone, spider mites showed no preference (main effect of the type of test: *F*_*2,63*_ = 1.01, *P* = 0.37; model 11; Figure 4A). Indeed, the time spent by females on each side of the arena was the same, irrespective of the type of choice presented to the mites (no droplet us. LB: t(1) = -1.54, *P* = 0.13; *E. coli* us. LB: t(1) = 0.06, *P* = 0.95; *P. putida vs.* LB: t(1) = 0.25, *P* = 0.81). Moreover, in all types of tests performed with the Y-maze olfactometer (main effect of type of test: *X*^*2*^_*3*_ = 3.86, *P* = 0.28; model 12) mites did not show any preference for the arm of the olfactometer free from bacteria (difference to 0.5 for *E. coli* vs LB on cotton: z=-1.12, *P* = 0.27; *P. putida* vs LB on cotton: z=-0.89, *P* = 0.37; *E. coli* vs LB on plant: z=0.22, *P* = 0.82; *P. putida* vs LB on plant: z=1.18, *P* = 0.18; Figure 4B). Thus, we found no evidence for the effect of volatile cues on spider-mite avoidance behavior toward bacteria.

**Figure 4.**
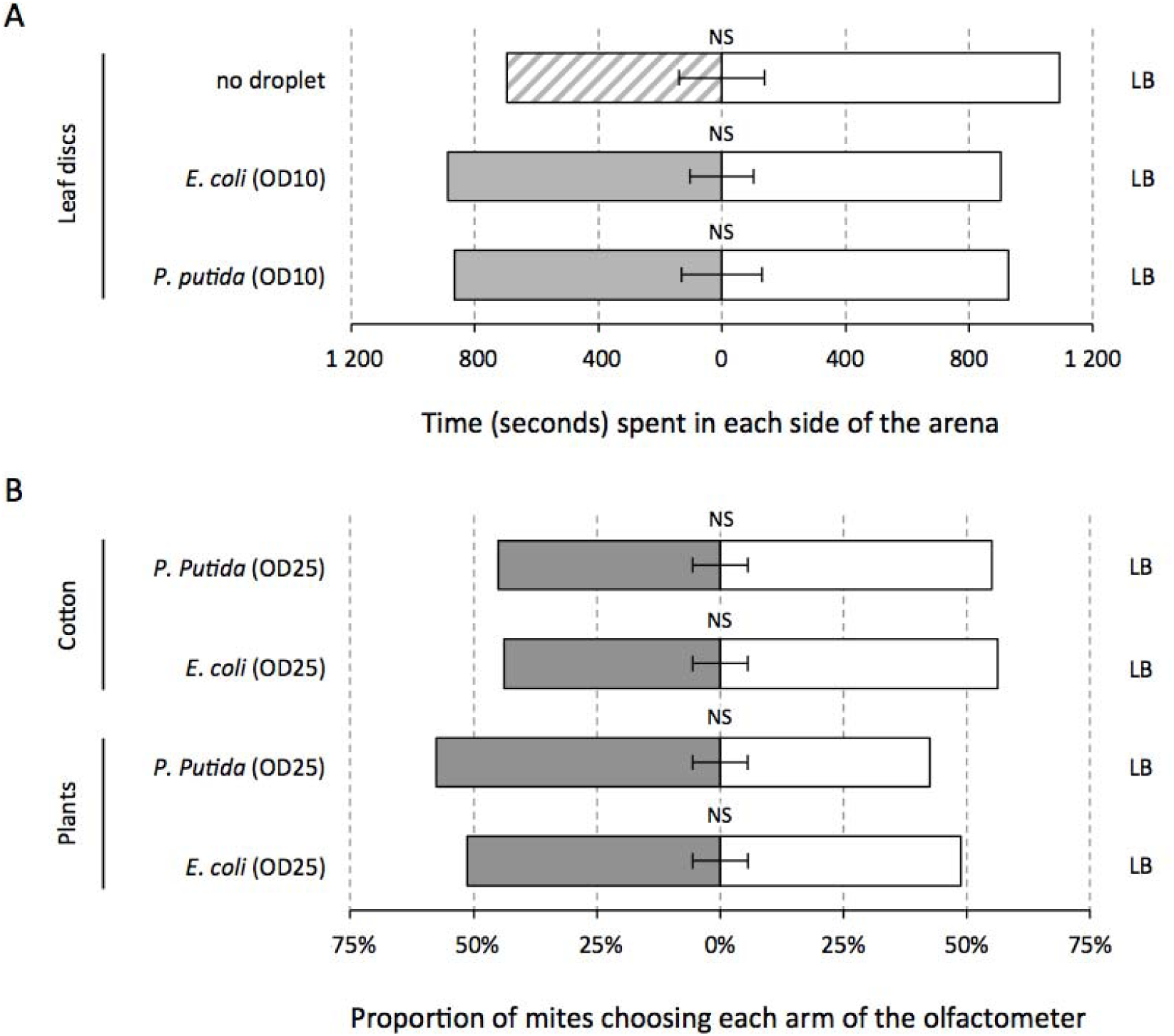
Effect of volatile cues on spider-mite avoidance behaviour. Bars represent mean (± s.e.) time spent by females close to a leaf disc free with either no droplet, or a droplet containing either LB or bacteria (*E. coli* or *P. putida)* at ODIO (A), and the proportion of females choosing an arm of the olfactometer containing either bacteria (E *coli* or *P. putida)* at OD25 or LB, on cotton or on entire bean plants (B). NS indicates that no significant differences have been found at the 5% level. White bars: LB; Light grey bars: OD1; Medium grey bars: ODIO; Dark grey bars: OD25; Dashed bars: absence of droplet.

## DISCUSSION

In this study, we found that spider mites die upon infection with *Escherichia coli* and *Pseudomonas putida* both by spraying and ingestion. Moreover, we show that they avoid contaminated food and that odours are probably not involved in such discrimination.

In a previous study, we have shown that mites do not mount an effective immune response towards systemic bacterial infections (Santos-Matos et al. 2017). This result was intriguing, as mortality upon infection was very high, indicating that bacteria impose severe fitness costs to spider mites, which should select for traits that allow avoiding such costs. Hence, we here tested for the generality of these costs for other putative infection routes, and for the presence of other potential defence mechanisms, namely behavioural. We found that spraying mites with bacteria also led to significant mortality, which suggests that bacteria gained access to the body cavity of the mites. This may have occurred by ingestion (as the sprayed bacteria remain in the leaves, hence mites may feed upon them), but most likely also through the cuticle (e.g. via spiracles or tracheae). Both cuticle and gut epithelium may thus be permeable to bacteria! infections. We then tested whether ingestion alone would be sufficient to induce severe mortality. We found this to be case. However, mortality levels induced by feeding alone were less severe than those observed with spraying (see Figures 1A and C and 2A and C), which can be due to differences between protocols, but also to additional mortality induced by bacteria penetrating the cuticle. Indeed, spraying may lead to a higher amount of bacteria per mite than feeding on contaminated food. Testing this hypothesis would, however, require data on the bacterial loads per mite after infection. Nevertheless, the fact that injection with *E. coli* led to 100% mortality even at the lowest dose tested (Santos-Matos et al. 2017) whereas, in this study, some mites survived following bacterial exposure through other routes, suggests that cuticular and epithelial barriers confer some degree of protection. Importantly, these barriers may be much more effective in nature than in our experimental conditions where the amount of bacteria used is extremely high. However, some degree of permeability and ensuing exposure to bacterial infections are likely to occur in nature and, as discussed below, may select for spider-mite behavioural strategies to avoid exposure to pathogens.

We observed different responses in oviposition upon infection. Overall, oral infection of spider mites with *E. coli* resulted in decreased oviposition rate, whereas spraying led to an increase of this trait. Conversely, oral infection with *P. putida* resulted in increased oviposition whereas spraying led to a reduction of this trait. An interesting pattern thus seems to emerge between virulence and oviposition, whereby low virulence entails lower fecundity and extremely virulent infections induce the opposite response (i.e. increased oviposition). Often, infection entails a cost in terms of oviposition reduction, as observed in many species, such as mosquitos (Pigeault et al. 2015; Scholte et al. 2006; Zele et al. 2018) or sticklebacks (Heins 2012). However, some hosts are able to compensate infection-driven fitness costs by changing the timing of their reproductive efforts (i.e. ‘fecundity compensation’; Parker et al. 2011; Vezilier et al. 2015). For example, *Daphnia magna* produces more offspring early in life when exposed to the microsporidian parasite *Glugoides intestinalis* (Chadwick and Little 2005). Our data suggest that the effects of bacterial infection on oviposition include both effects described above, with cost or compensation being contingent upon bacterial species and infection mode, as shown in other systems (Martins et al. 2013). More information on infection dynamics in spider mites, the relationship between total reproductive output and longevity, as well as how it affects other life-history traits, such as egg hatching, will help clarify this issue.

Independently of the route of infection tested, *E. coli* was less pathogenic than *P. putida.* Although a same bacteria! density measured at OD_600_ does not correspond to the same number of bacteria for each species, the increased pathogenicity of *P. putida* cannot be due to an increased concentration compared to *E. coli* (i.e. a lower number of *P. putida* than off. *coli* result in the same OD). Moreover, this result is consistent with previous work using systemic infection in *Drosophila* (Martins et al. 2013), and thus generalizes the virulence hierarchy between these bacteria beyond the infection route. *E. coli* is a particularly mild pathogen compared to *P. putida.*

We found that the avoidance response regarding *P. putida* ingestion was stronger than that observed for *E coli.* Possibly, spider mites detect *E. coli* to a lesser extent than they detect *P. putida,* and/or the avoidance level depends on the magnitude of the threat. In our study, mites showed higher avoidance in feeding on *P. putida,* which is more pathogenic than *E. coli.* This would suggest both that bacterial avoidance is costly (see Buck et al. 2018) and consequently selected only for highly pathogenic bacteria, and that spider mites are able to discriminate between different degrees of pathogenicity. Within bacterial species, mites avoided more often *E. coli* at high concentrations and less at low concentrations, but were equally affected by all concentrations tested. This may entail that, under natural conditions, low *E. coli* concentrations will lead to higher mortality within populations. However, this was not the case for *P. putida,* as high concentrations of this bacterium induced a higher mortality and were more avoided by spider mites than low concentrations. The avoidance to feed on contaminated food is not necessarily triggered by an avoidance of infection per se. Indeed, mites may also use bacteria as a proxy for low-quality food, such as rotten food (Janzen 1977). Conversely, in absence of a direct detection of pathogens, hosts may use rotten food as a cue that is reliably associated with pathogens to avoid becoming infected (see Buck et al. 2018). Thus, the feeding avoidance observed in our experiment may reflect a more general response against infection by diverse pathogens.

We found that avoidance was not triggered by olfactory cues, either directly produced by the bacteria or resulting from their interaction with the plants. We cannot exclude the possibility that these results are due to limitations in our experimental setup. However, both set-ups have been previously used to show the ability of spider mites to detect olfactory cues, albeit in other contexts (Fouks and Lattorff 2011; Pallini et al. 1997; Parker et al. 2010; Rodrigues et al. 2017; Rondot and Reineke 2017; Tasin et al. 2012). Nevertheless, our results suggest that avoidance happens locally, in the plant, and not at long distance, and are consistent with spider mites not perceiving bacteria odours but, instead, avoiding rotten food after tasting. Indeed, gustatory avoidance has been previously shown for other arthropods, such as *Drosophila* avoiding bacterial LPS through the activation of the protein TRPA (Soldano et al. 2016).

Gustatory avoidance is likely to have strong effects locally, both in terms of host mortality and parasite transmission. The survival of spider mites is drastically reduced when bacteria penetrate their body as they have a deficient immune response upon infection (Santos-Matos et al. 2017). Moreover, bacteria proliferate inside their body (Santos-Matos et al. 2017), which can increase the likelihood of transmission (Anderson and May 1979). Consequently, individuals with reduced avoidance are more likely to be exposed and, thus, to die upon bacterial infection. Natural populations of spider mites are frequently at high-density of individuals, which favours pathogen transmission (Grbic et al. 2011). If individuals do not avoid pathogens and bacteria proliferate within hosts, this might lead to an epidemic and, in an extreme case, to population extinction.

Our results show that spider mites, while not mounting an effective immune response upon infection by bacteria, have alternative defensive strategies. These include the cuticle and gut barriers of the body, apparent fecundity compensation following infection, and behavioural avoidance of contaminated food sources. Therefore, these data highlights the importance of investigating different host defensive strategies in order to fully analyse the outcome of pathogen infection. Moreover, our results highlight the need of further research to determine the relevance of such findings in nature.

## ACKNOWLEDGEMENTS

We thank Miodrag Grbic for the “bubble machine” and José Feijó for the air pump, both of which were necessary to make the parafilm bubbles (feeding experiments). We thank Arne Jansen and Luísa Vasconcelos for the material needed to make the Y olfactometer, and Francisco Dionísio for providing access to his laboratory, material and reagents for bacterial work. We also thank members of the Sucena and Magalhães labs for useful discussions and suggestions. This work was funded by an FCT-Tubitak agreement (FCT-TUBITAK/0001/2014) to SM and Ibrahim Cakmak. FZ was funded through an FCT Post-Doc fellowship (SFRH/BPD/125020/2016). Funding agencies did not participate in the design or analysis of experiments. We declare that we do not have any conflict of interest.

**AUTHORS’ CONTRIBUTIONS**
Experimental conception and design: FZ, GM, ES, SM; acquisition of data: GM, AF, CE, CP, TL; statistical analyses: FZ; paper writing: FZ, GM, SM, with input from all authors. Funding: ES, SM. All authors have read and approved the final version of the manuscript.

